# Interactions retain the co-phylogenetic matching that communities lost

**DOI:** 10.1101/033050

**Authors:** Timothée Poisot, Daniel B. Stouffer

**Affiliations:** Centre for Integrative Ecology, School of Biological Sciences, University of Canterbury, Christchurch, New Zealand; Université de Montréal, Département de Sciences Biologiques, Montréal, Canada; Québec Centre for Biodiversity Sciences, Montréal, Canada

**Keywords:** species interactions, host-parasites, phylogenetic congruence, network cophylogeny, community phylogenetics

## Abstract

Both species and their interactions are affected by changes that occur at evolutionary time-scales, and these changes shape both ecological communities and their phylogenetic structure. That said, extant ecological community structure is contingent upon random chance, environmental filters, and local effects. It is therefore unclear how much ecological signal local communities should retain. Here we show that, in a host–parasite system where species interactions vary substantially over a continental gradient, the ecological significance of individual interactions is maintained across different scales. Notably, this occurs despite the fact that observed community variation at the local scale frequently tends to weaken or remove community-wide phylogenetic signal. When considered in terms of the interplay between community ecology and coevolutionary theory, our results demonstrate that individual interactions are capable and indeed likely to show a consistent signature of past evolutionary history even when woven into communities that do not.

---

Ecological interactions often exert important selective pressures on the species involved. For example, the phenologies of lodgepole pines and red crossbills respond spatially to the presence of squirrels (Benkman et al. 2003). Likewise, palm species undergo changes in seed morphology in response to the extinction of birds dispersing their seeds (Galetti et al. 2013; Johnson et al. 2017). Interactions can be lost, too, when phenologies of the species involved shift (Rafferty et al. 2015). Interactions are, in fact, so important that the existence of a species has been inferred by the fact that another species bore traits that matched no other known species: Kritsky (1991) relates the discovery of the moth *Xanthopan morganii*, with a proboscis famously over a foot long, which Darwin predicted would exist based solely on the phenology of local plant *Angraecum sesquipedale*. In addition, interactions and the emergent structures they define are distributed in similar ways across communities at both large or small scales (Jordano et al. 2003). Together, these observations suggest that much ecological structure could be the end result of (co)evolutionary dynamics between species (Eklöf et al. 2012; Stouffer et al. 2012). Unfortunately, although the evolutionary dynamics of pairs of interacting species have been well described at macro-evolutionary (Van Valen 1973) and micro-evolutionary (Gandon et al. 2008) timescales, most attempts to understand how they cascade up to the levels of diversity of both species and interactions found within empirical communities have been inconclusive (Hembry et al. 2014). This suggests that these well-described mechanisms may not confer substantial predictive power when examined at scales of organization larger than the pairwise interaction.

Historically, the evidence for shared evolutionary history in taxonomically diverse communities relied on the quantification of the degree of matching between the phylogenies of two sets of interacting organisms, accounting for the distributions of interactions across the phylogeny (Legendre et al. 2002). This notion builds on the century-old idea that extant species interact in a way similar to the way their ancestors did (Fahrenholz 1913; Guimarães Jr et al. 2011; Nuismer et al. 2013). Note that testing these assumptions is related to, but markedly more restrictive than, testing for phylogenetic conservatism of species’ interactions (Rezende et al. 2007; Eklöf et al. 2012). This is because of additional, higher-order constraints related to the shape of both trees at *all* depths (Cavender-Bares et al. 2009; Mouquet et al. 2012), because ancestral evolutionary innovations have a high phylogenetic inertia, and they carry forward to extant taxa (Desdevises et al. 2003; Diniz-Filho & Bini 2008; Vale & Little 2010). In a way, the true measure of phylogenetic signal of interactions should depend not only on how they are conserved within the tree of the species establishing them (*e.g.* parasites or pollinators), but also how these interactions at matched to the tree of the species receiving them (*e.g.* hosts or plants). This is true whether or not the species complex are coevolving: in fact, neutral interactions can yield perfectly matching co-phylogenies (Poisot 2015). Consequently, many of the systems that have been described as exhibiting significant phylogenetic structure of interactions ultimately deviate from this last constraint, and this can occur for a variety of factors that stem from how other species evolved and established, lost, or maintained interactions throughout their joint evolutionary history (in addition to the fact that species may have different impacts on one another, *i.e.* the effect of any interaction on the community-level co-phylogenetic signal is not expected to be equal). Nonetheless, detecting matching phylogenies for interacting clades indicates that their shared evolutionary history is long standing and is therefore suggestive that their extant ecological structure is an outcome of ancestral constraints and/or co-adaptation (Nuismer & Harmon 2015).

It is important to note further that discovering matching phylogenies does not mean that coevolutionary dynamics—*sensu* Thompson (1999)—took place at any time. In fact, coevolution is not expected to necessarily result in matching phylogenies nor are matching phylogenies only produced through coevolution (Poisot 2015). It follows that community-level measures of phylogenetic signal, while they *do* quantify how closely interactions are a product of phylogeny, do not allow us to draw conclusions on coevolution. Nevertheless, interaction-level measures are useful in that, when expressed as the contribution of interactions to the overall signal, they allow us to *compare* the importance of interactions across replicated communities. Communities from the same regional pool vary because (i) the local species pool is at best a subset of the regional species pool and (ii) the local interactions are at best a subset of the interactions in the regional community (Poisot et al. 2015). This implies that (i) the phylogenetic signal in the regional pool will be different from the signal in the local communities, and (ii) the phylogenetic signal across local communities will differ. Species sampling and variability of interactions, however, do no predict (i) how the phylogenetic signal of pairwise interactions is kept or lost at the scale of the whole community nor (ii) whether or not this variability is related to changes in the amount of phylogenetic signal that can be detected locally.

In this manuscript, we analyze a large dataset of over 300 species of mammalian hosts and their ectoparasites, sampled throughout Eurasia, for which phylogenetic relationships are known. Using a Procrustean approach to quantify the strength of co-phylogenetic matching of interactions between host and parasite trees (Balbuena et al. 2013), we show that locally sampled communities rarely show strong matching despite the fact that the overall system does at the continental scale. We then provide evidence to support the conclusion that the amount of phylogenetic matching within a local community is predictable based on the importance of interactions in the regional network. We finally show that the contribution of specific interactions to phylogenetic matching is invariant across scales, and is unrelated to their tendency to vary across space. The lack of co-phylogenetic structure in local communities suggests that, while interactions are undeniably important for community assembly, they might be less so than abiotic factors.

## 1 METHODS

### 1.1 Data source and pre-treatment

We use data on observations of interactions between 121 species of rodents and 205 species of parasitic fleas in 51 locations across Europe (Krasnov et al. 2012b) to build 51 species-species interaction networks. Interactions were measured by combing rodents for fleas, a method that gives high quality data as it has a high power of detection. The dataset also includes phylogenies for the hosts and the parasites. Previous analyses revealed that this dataset shows significant co-phylogenetic matching at the continental level (Krasnov et al. 2012a). Importantly, it also provides spatial replication and variability (Canard et al. 2014) at a scale large enough to capture macro-ecological processes. This dataset is thus uniquely suited for our analysis as it represents a thorough spatial and taxonomic sampling of a paradigmatic system in which interspecific interactions are thought to be driven by macro-evolution and co-speciation events (Combes 2001; Verneau et al. 2009).

The original dataset gives quantitative interaction strengths (expressed as an averaged number of parasites per species per host). In this system, quantitative interaction strengths were previously shown to be affected to a very high degree by local variations in abundance across sampling locations (Canard et al. 2014), and it therefore seems unlikely that they reflect macro-ecological processes. Therefore, to account for differential sampling effort—which cannot readily be quantified—and across site variations in abundance—which do not pertain to macro-evolutionary processes—we only study the networks’ bipartite incidence matrices (that is, presence and absence of infection of hosts by the parasites).

### 1.2 Spatial scales and interaction spatial consistency

Noting that variation of interactions across locations—which can be caused by local ecological mechanisms as opposed to reflecting evolutionary dynamics—can decrease congruence, we analyze the data at three different levels which we will refer to as continental, regional, and local. Notably, the continental level summarizes the complete dataset whereas both the regional and local levels are location-specific scales.

*Continental* interaction data consists of the aggregated “metanetwork” which includes all documented interactions between species from the regional species pool (Poisot et al. 2012). For every location, we further define two scales of analysis.

First, *regional* interaction data accounts for different species composition across sites: the species that have been observed locally interact as they would do in the *continental* network, *i.e.* if interactions did not vary across space. This allows testing whether sampling from the regional species pool affects co-phylogenetic matching, regardless of the distribution of interactions. Within each site, the regional scale is given by the subset of the metanetwork formed by the locally present species (properly speaking, the induced subgraph of the metanetwork induced from the nodes of the local network). Hence the regional networks are always a perfect subset of the continental network, and do not reflect whether species were actually observed to interact locally or not, but whether they *can* interact at all. This *regional* network is thus a baseline estimate derived from interactions within the species pool and measures the effect of species sampling on co-phylogenetic matching.

Second, *local* interaction data are the actual observations at this location: the identity of the species present, and the way they interact. In addition to capturing the dissimilarity of species composition across sites, this allows measuring the effect of interaction turnover across space. The local and regional networks always include the same species, but the local network has only a subset (or, at most, an exact match) of the interactions in the regional network.

We finally define the spatial consistency of every interaction as the proportion of sites in which the two co-occurring species interact with each other, or simply

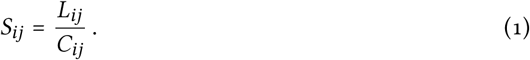

The spatial consistency of an interaction *S_ij_* between species i and j is therefore the ratio between the the number of locations in which they were observed to interact (*L_ij_*) and the number of locations in which both were observed to be present (*C_ij_*). Because *L_ij_* ∈ [0, *C_ij_*], this measure takes values in [0, 1] where larger values reflect high spatial consistency. Note that although they are reported as 0 (*i.e.* having no interactions), we actually have no information about species pairs that have never co-occurred; this is a common, but hard-to-correct-for, feature of spatially replicated datasets in which species occurrence varies (Morales-Castilla et al. 2015). Therefore, the only values of *S_ij_* can be properly estimated are those for species pairs that have been observed to *co-occur* at least once.

### 1.3 Quantifying co-phylogenetic matching

We quantify the strength of co-phylogenetic matching in terms of the degree of matching between host and parasite phylogenies given knowledge of extant species interactions. We do so using the *PACo* method (Balbuena et al. 2013), which is robust to variations in both number of species and interactions. *PACo* provides measures of both the network-level congruence (*i.e.*, is there phylogenetic signal in the species interactions across the entire network?) and the interaction-level signal (*i.e.*, what is the contribution of each interaction to the overall signal?). Because interaction-level measures provided by *PACo* operate like residuals, larger values of this metric reflect *low* contributions to co-phylogenetic matching. Likewise, interactions that contribute strongly to phylogenetic congruence have smaller *PACo* values. Importantly, and in contrast to previous methods such as *ParaFit* (Legendre et al. 2002), *PACo* also can be used to meaningfully quantify the contribution of every interaction to the network-level signal even in cases where the entire network shows no significant phylogenetic signal.

All values returned by *PACo* are tested for deviation from a random expectation (Hutchinson et al. 2017), and we generated those random expectations by applying permutations to the species interaction networks. Specifically, we applied permutations that maintained the number of parasites for each host and the number of hosts for each parasite. This has the effect of measuring whether re-distributing interactions between tree tips would give rise to the same value. We always compared the observed value to the randomized distribution using a two-tailed statistic; thus, a significant value indicates that the observed value is unlikely to have been observed by chance, without pre-specifying whether or not it is larger or smaller than expected.

In PACo, the effective sample size is the number of interactions in the network, and our interpretation of PACo's output must account for this. This is not an issue for permutation tests, since they evaluate the significance of the co-phylogenetic signal by permutations of each network, the power of each test varies but the test statistics can be compared. To ensure that values of the interaction contribution to co-phylogenetic signal are comparable, we normalized them network-wise by dividing them by the maximal value of the sum of squares in PACo. While the raw values returned by PACo are not meaningfully comparable between networks, the corrected values presented here are.

As required by *PACo*, the phylogenetic trees for hosts and parasites were rendered ultrametric (*i.e.*, all species are at the same distance from the root). This has the consequence of losing the temporal component of the tree (which was not available for the parasites in the original dataset), but standardizes phylogenetic distances in a way that satisfies *PACo*’s requirements. Moreover, this introduces the hypothesis that the common ancestor to the parasites was able to infect the common ancestor of the host. Note that the same procedure was applied in the original publication based on these phylogenetic data (Krasnov et al. 2012a).

## 2 RESULTS AND DISCUSSION

Splitting the datasets at the continental, regional, and local levels delineates clear quantitative predictions. At the regional scale, one can expect community assembly to promote the co-occurrence of evolutionarily linked species pairs – *i.e.*, a host and a parasite from lineages that interact will tend to co-occur more often because the parasites are filtered to be present in sites where they can find hosts. Under this situation, we expect that regional networks will have a high degree of phylogenetic matching (because they account for the information on potential species interactions); we do in addition expect that their phylogenetic signal will be larger than what is found in the continental network, since the latter represents a somewhat artifactual agglomeration of species pairs that do not co-occur. The opposite situation (a relatively lower phylogenetic matching) would therefore be suggestive of a weaker selection for the co-occurrence of evolutionarily tied species pairs.

At the local scale, if interactions between species at matching phylogenetic positions are conserved, we would expect both a similar or higher level of phylogenetic matching between the local and the regional scale, and a positive relationship between the frequency of interaction and its overall importance for phylogenetic matching (interactions with a strong phylogenetic signal happen more often). On the contrary, if local assembly proceeds largely independently from the coevolutionary history, the relative level of phylogenetic matching in local networks should be the same as in the regional networks (through a sampling effect from the distribution of interaction-level contribution to co-phylogenetic matching), but the frequency of interactions should bear no relationship to their importance in overall matching.

### 2.1 Local and regional scale networks show no co-phylogenetic matching

As host-macroparasite interactions are hypothesized to be ecologically constrained due to their being evolutionary conserved (Combes 2001), the congruence observed at the continental level sets the baseline for what would be expected in local communities. Of course, if ecological mechanisms (such as filtering) reduce co-phylogenetic matching, we should detect this signal at the continental scale but not locally. Out of 51 sites, our *PACo* analysis indicates that 35 show no signal of co-phylogenetic matching at all, 11 show significant co-phylogenetic matching when using the regional interactions, and 12 show significant co-phylogenetic matching using the local interactions (see *Supp. Mat. 1* for network-level significance values; table 1; fig. 1). These results support the idea that macro-evolutionary processes, such as co-diversification, can have consequences at the macro-ecological level but may not in fact be detectable at finer spatial scales.

**Table 1.** Contingency table showing the number of networks with (non-)significant local and regional co-phylogenetic signal.

|  | Regional non-significant | Regional significant |
| --- | --- | --- |
| Local non-significant | 35 | 4 |
| Local significant | 5 | 7 |

**Figure 1.**
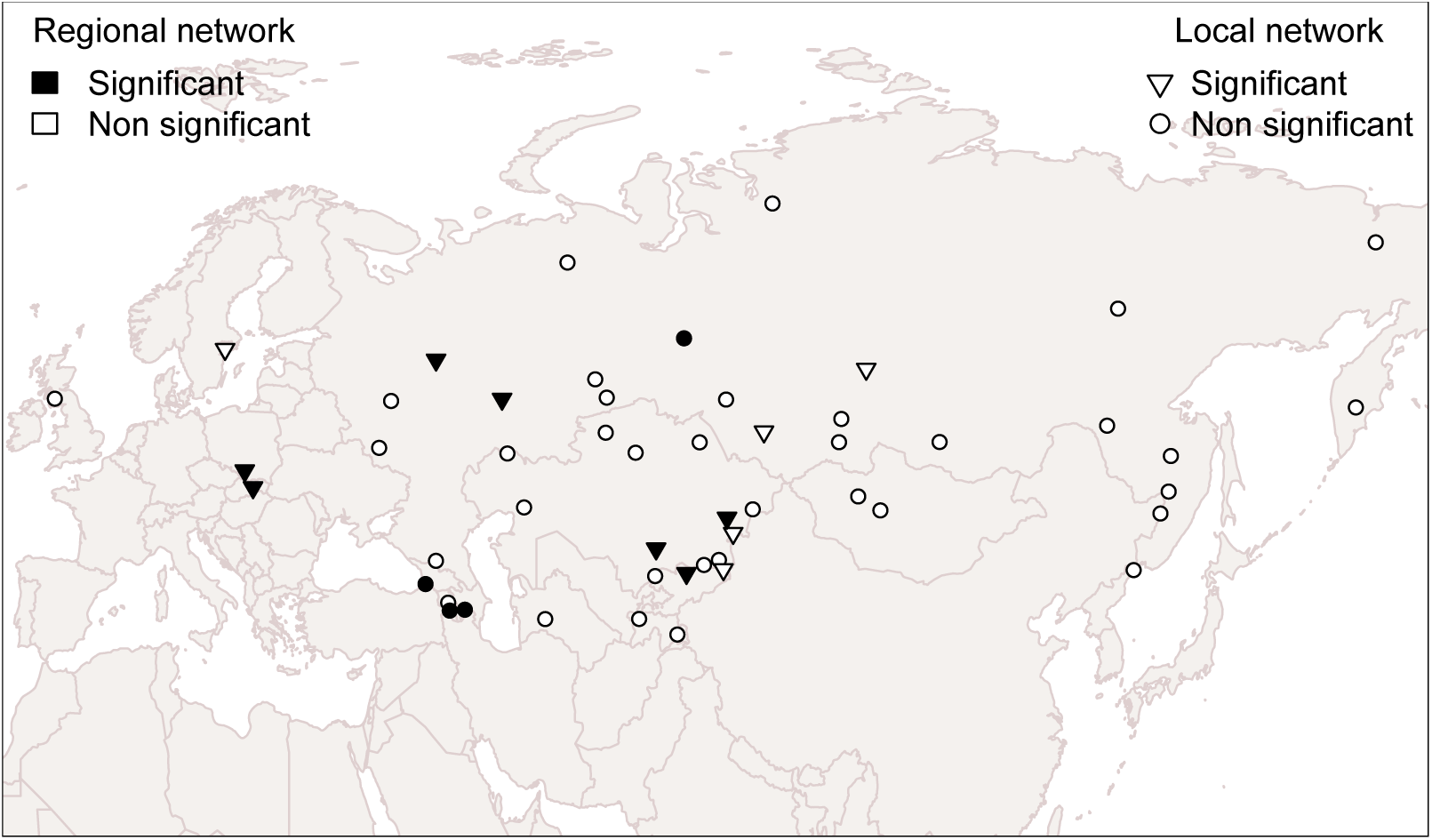
Spatial distribution of co-phylogenetic matching across the 51 sites. For each location, we indicate whether or not the structure of regional and local interaction networks is consistent with phylogenetic congruence. Locations where the local network shows a significant co-phylogenetic structure are marked with triangles. Locaitions where the regional network shows a significant co-phylogenetic structure are in black.

### 2.2 Local and regional scale networks have the same relative co-phylogenetic matching

When we compared the relative degree of co-phylogenetic matching in the local and regional communities (fig. 2), we see that the relationship between the two is approximately linear (95% confidence interval for the correlation coefficient 0.914–0.971). This fits with the hypothesis of local networks being assembled by a random sampling from regional networks: if interactions between species at matching positions in both trees are maintained by the same set of drivers, then this should be reflected in the local networks by a higher degree of co-phylogenetic matching.

**Figure 2.**
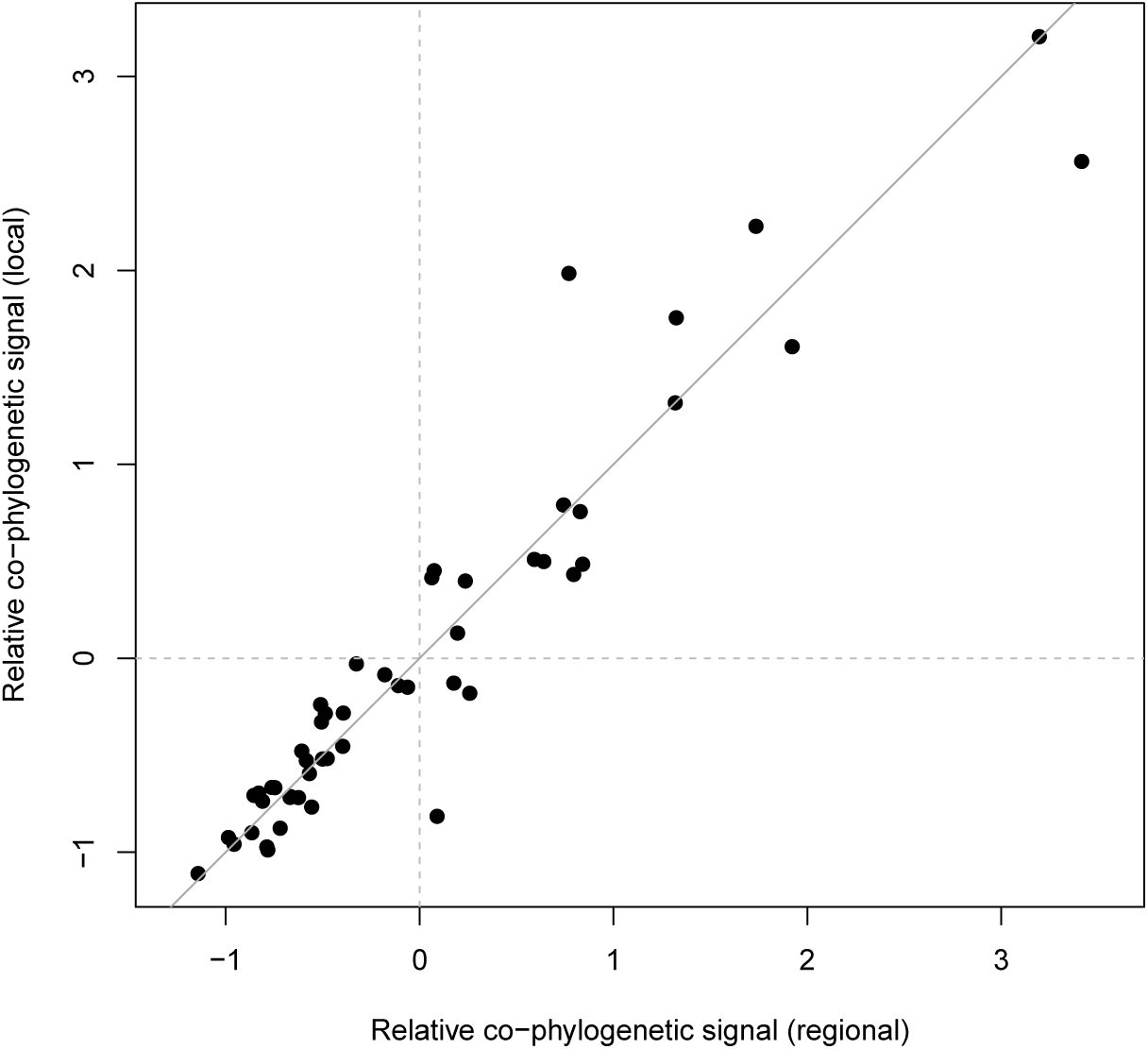
The regional and local networks show the same relative amount of co-phylogenetic matching. The values presented are the z-scores of the PACo statistic for the entire network, with the 1:1 relationship indicated by the solid line.

### 2.3 Co-phylogenetic matching is predicted by the contribution of interactions

On the other hand, system-level differences say little about the behavior of individual interactions. Despite the fact most coevolutionary mechanisms act at the interaction level (Thompson 1999), most *measures* of it are expressed at the community level. We observe here that networks with interactions that are important for co-phylogenetic matching at the continental scale are also important for co-phylogenetic matching at the local and regional scales as well (*ρ* = 0.95; fig. 3A). Intriguingly, we also find that the distribution of individual interactions’ contributions to co-phylogenetic matching is strongly conserved, regardless of the scale at which the interactions are quantified (fig. 3B). Because interactions differ between each other in terms of their total contribution to co-phylogenetic matching, this implies that their distribution across networks (*i.e.* whether the local network contains a sample of strongly contributing, or weakly contributing, interactions) is what actually drives differences in overall co-phylogenetic matching. As such, network-level co-phylogenetic matching emerges directly from the properties of interactions and is not a property of the network itself.

**Figure 3.**
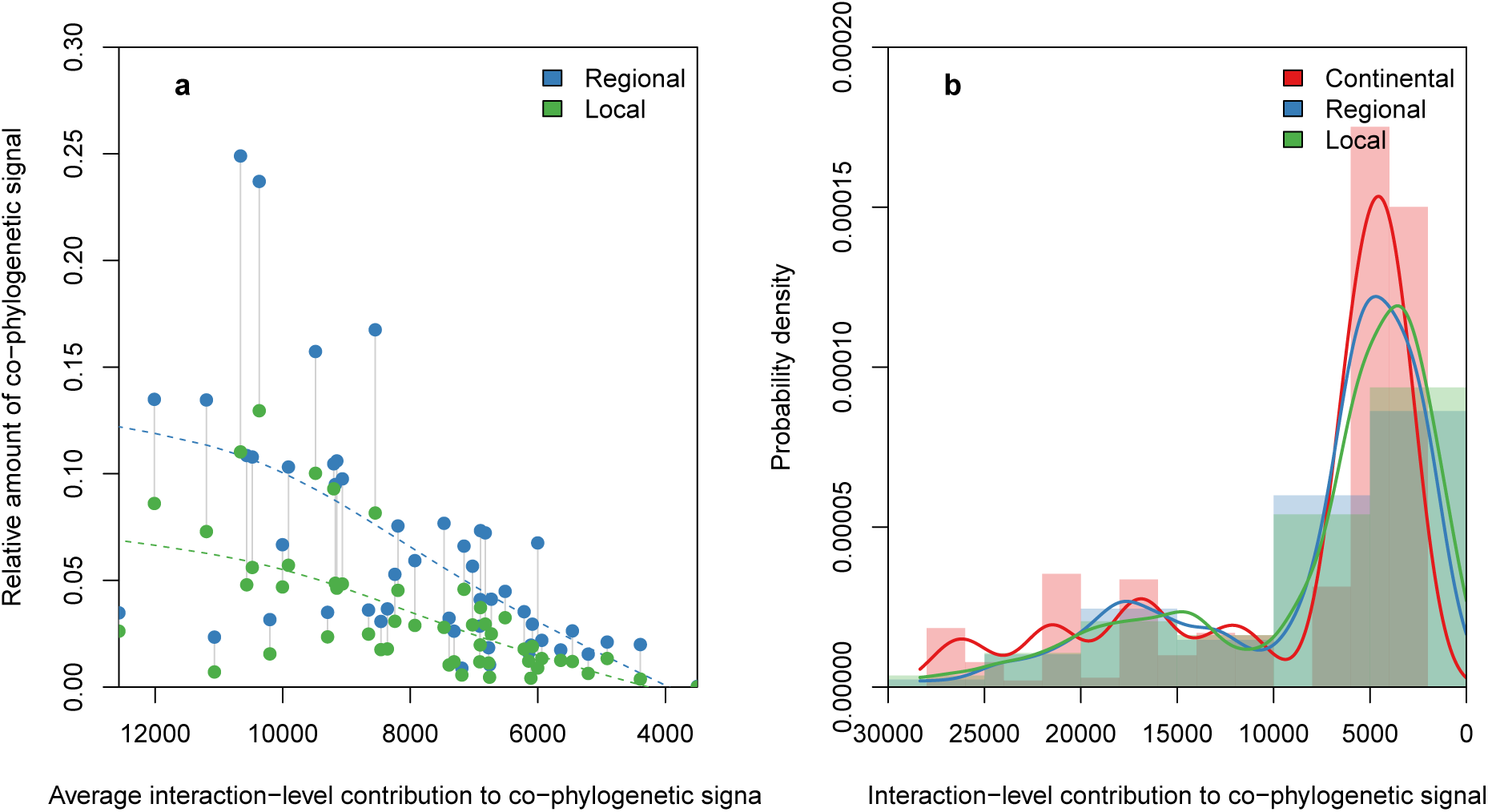
Distribution of co-phylogenetic matching at the network and interaction levels. **a**, Networks that have lower co-phylogenetic matching at the local or regional level are composed of interactions that on average contribute little to co-phylogenetic matching at the continental scale. Co-phylogenetic matching is presented relatively to the continental level co-phylogenetic matching. Dashed lines are a cubic smoothing spline, and the two levels of the same networks are linked by solid grey lines. **b**, Overall, interactions observed at the local, regional, and continental scale have roughly equivalent contributions to co-phylogenetic matching. Probability density was smoothed using a Gaussian kernel density estimator. Raw probability densities are shown as semi-transparent bars. Interaction-level contributions are unitless, and lower values represent stronger signal.

### 2.4 Interactions contributing to co-phylogenetic matching are marginally more spatially consistent

Beyond their contribution to co-phylogenetic matching, interactions also ultimately differ in how frequently they vary when the species involved co-occur (Carstensen et al. 2014; Olito & Fox 2015; Trøjelsgaard et al. 2015). This can happen, for example, when one of the partners is able to forage for optimal resources (Betts et al. 2015). Once more, the literature on host-parasite interactions assumes that the reason why some interactions are more frequent is because they reflect a significant past history of coevolution (Guimaraes et al. 2007; Nieberding et al. 2010); that is, the ecological constraints emerge from evolutionary conservatism. Using a weighted Pearson's correlation between the interaction frequency, interaction contribution to co-phylogenetic matching, and the number of observations of each interaction as the weight, we observe that this is marginally true (*ρ* ≈ −0.11. *t* ≈ −5.09 with weights; *ρ* ≈ −0.10, *t* ≈ −4.6 without; both significant at *α* = 0.05; fig. 4). Recall that the *negative* correlation here arises from the fact that high interaction-level values in PACo means *low* contribution to co-phylogenetic signal. Nevertheless, the significance of this result ought to be tempered by the fact that the *R*^2^ of both regressions is close to 0.01. Consequently, the association between spatial consistency and contribution to co-phylogenetic signal, while statistically significant, explains so little variance of either quantities that it is likely of negligible biological importance. This implies that the spatial consistency of an interaction does not necessarily reflect its evolutionary past, but rather (possibly) extant ecological processes.

**Figure 4.**
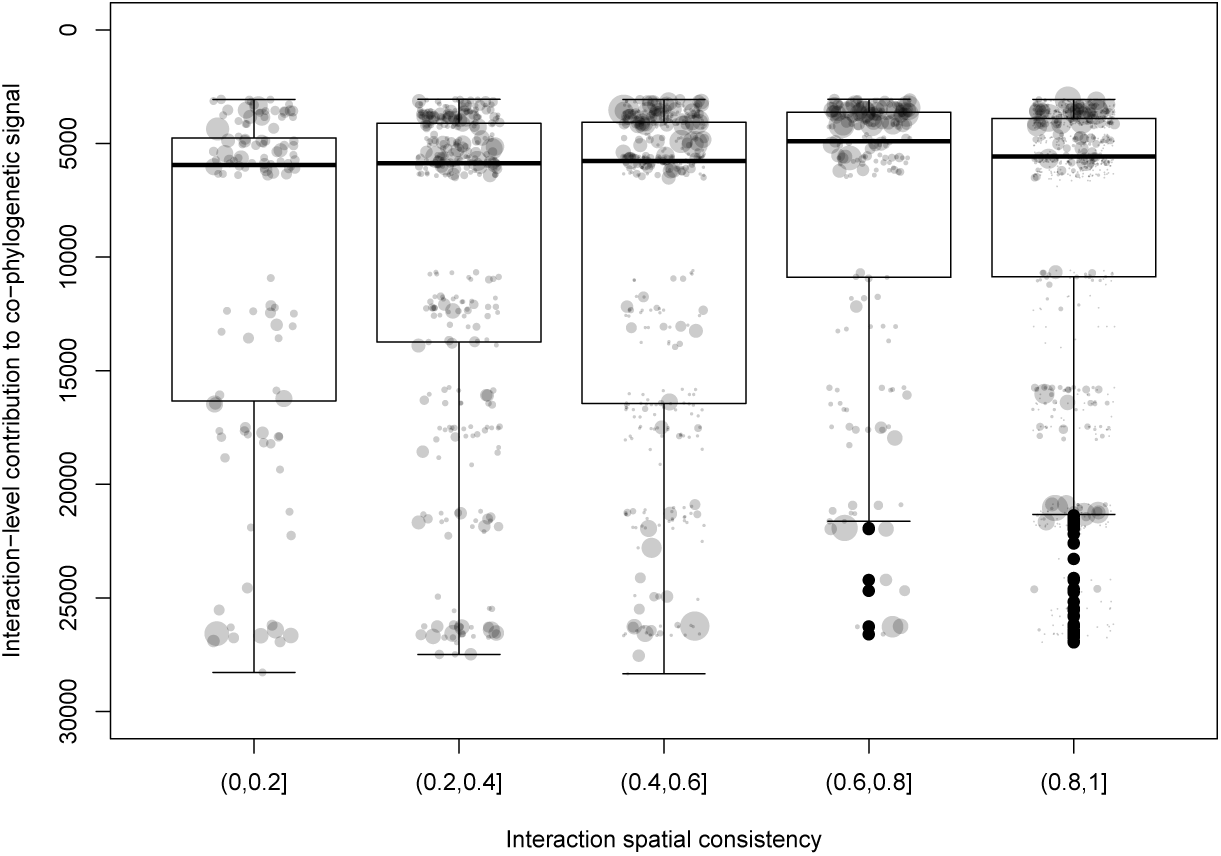
Spatial consistency of an interaction and its contribution to co-phylogenetic matching. Note that because *PACo* gives low scores to interactions with a strong contribution to co-phylogenetic matching, the y-axis is reversed. Spatial consistency is defined as the probability of observing an interaction between two species given that they were observed to co-occur (the size of the point is proportional to the number of times an interaction was observed). Although statistically significant, there was no biologically meaningful relationship between spatial consistency and an interaction's importance for co-phylogenetic matching in the continental network (*R*^2^ ≈ 0.01, *ρ* = −0.1, *p* ≤ 10^−5^).

### 2.5 The contribution of interactions to co-phylogenetic matching is consistent across scales

Ultimately, co-phylogenetic matching varies across scale because of the simultaneous variation of species’ interactions *and* communities’ phylogenetic tree structure. In a system characterized by substantial turnover, we would expect the contribution of each separate interaction to differ across scales as well. Instead, we observe here that interactions that contribute strongly to co-phylogenetic matching at the continental scale *also* show a significant tendency to contribute strongly at the local (*p* < 0.05 for positive correlations in 48 out of 51 networks) and regional (in 47 out of 51 networks), and this observation is independent of network-wide co-phylogenetic matching (fig. 5). Remarkably, this result implies that the remnants of co-phylogenetic inertia are still locally detectable in *individual interactions* even though shared evolutionary history regularly fails to leave its imprint on most local networks.

**Figure 5.**
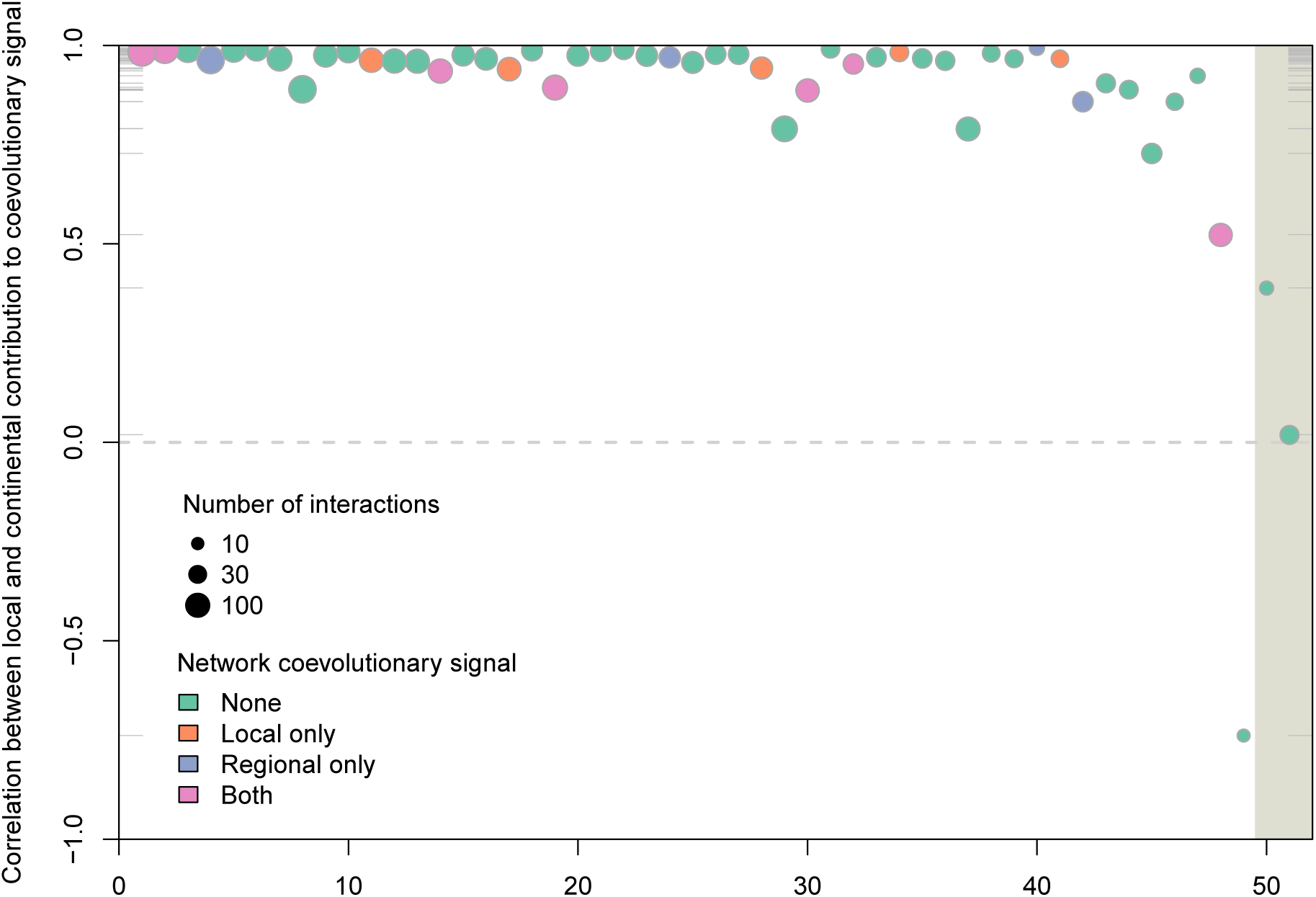
The contribution to co-phylogenetic matching of the interaction between two species is maintained across scales. For every site (ranked on the x-axis), we show the Pearson's correlation between interaction-level values of co-phylogenetic matching in the continental network and the same in the local network. The size of each point is proportional to the size of the network, and correlations for all sites are significant at *α* = 0.05 except for those falling in the grey shaded area.

## 3 CONCLUSIONS

Overall, the results of our study demonstrate that there is a sizeable gap between our current understanding of host-parasite coevolution as the basis of multi-species interactions, its phylogenetic consequences, and their applicability to ecological questions. Our results suggest that, while the continental-scale system might show a strong signal of past coevolution through significantly matching phylogenies (which was also reported, through different analyses, by other studies of this system), the quasi-entirety of this signal is lost when species and their interactions are filtered to assemble local communities. That there is no further loss of signal from the regional to the local scale strongly suggests that the loss of signal from the continental to regional scale is due to species sampling in a manner that acts independently of the evolutionary history of species pairs. Because regional and local networks have the same species, the difference between them stems from the loss of some interactions locally. It would therefore seem that local species pools in this system are also driven by the interaction between abiotic conditions and species tolerance, in addition to potential species interactions. Taking a step back, this result suggests that, while a shared phylogenetic history is a strong structuring force at the scale of the species pool, its influence is overridden by other factors during species filtering and community assembly. It further highlights the need for future investigation of whether the importance of phylogenetic history in host-parasite assemblages decays at smaller spatial scales.

Recently, Coelho et al. (2017) studied the factors that shape phylogenetic signal in bipartite interactions between plants and mutualists. In contrast to our study, they used a Mantel test to compare the two trees, a method whose lack of robustness (Harmon & Glor 2010; Guillot & Rousset 2013) is what originally prompted the development of PACo (Balbuena et al. 2013) or ParaFit (Legendre et al. 2002). Setting aside the relative merits of the various methods, they showed that phylogenetic signal is expected to decrease when (1) dispersal probability increases (*i.e.* all locations share the same species) and (2) the probability of interactions is high (*i.e.* there is no loss of signal due to local interaction filtering). We argue here that the probability of interaction should not be viewed as a system-wide measure, but is better defined for each interaction separately (eq. 1). When this is the case, we report no relationship between the probability of interaction and its contribution to co-phylogenetic signal (fig. 4).

The lack of a relationship between an interaction's probability of occurring (which represents their ecological stability) and its contribution to the co-phylogenetic process (which can be a proxy of their evolutionary stability) draws into question the basic long-term stability of interactions. Specifically, the existence of interactions with strong co-phylogenetic contribution, but low probability, is puzzling. It is tempting to hypothesize that these interactions may have once been strong enough to drive (co-)speciation, but are now decaying, due either to the fact that the host-parasite pair has been disrupted (by additional speciation and extinction events in the hosts or parasites) or as a consequence of ecological processes (for example, changes in the environment modifying the strength of some interactions). Janz (2011) show that parasites can frequently acquire hosts that are not closely related to their initial host. This phenomenon, dubbed the “parasite paradox”, can be responsible for a large number of diversification events, as parasites will be introduced to a novel environment when exposed to a novel host. The acquisition of novel hosts proceeds primarily from ecological contact between distantly related species. This emphasizes the need to account for the spatial structure of species interactions in these studies – because all species do not co-occur at all places, the potential for ecological contact with novel hosts varies across space. Ideally, reconstruction of the ancestral range of the species can shed some light on whether acquisition of distantly related species happened, and where.

Local networks show little to no signal of co-phylogenetic matching, and the strength of co-phylogenetic matching that can be ascribed to the interactions between two species is a surprisingly poor predictor of how frequently they interact. In contrast to the frequent assumption that phylogenetic structure is a key driver of community structure (Cavender-Bares et al. 2009), these data reveal that this impact is actually minimal at ecologically relevant spatial scales, even in a system where phylogenetic structure would be expected to have a profoundly strong role. And yet, despite all the above, individual interactions are somehow able to maintain their co-phylogenetic matching even when the community they are woven into does not. Thinking more broadly, these discrepancies provide a roadmap for bridging the gap between our appreciation of the role of shared evolutionary history and its empirically measurable outcomes: network structure is the most parsimonious *mechanism* by which coevolution proceeds, not the imprint potential coevolution leaves on ecological communities (Ponisio & M'Gonigle 2017).

Finally, we would like to acknowledge some limitations in the data. Host-parasite evolution usually follows a number of canonical coevolutionary scenarios (Page 2003): true cospeciation, where the two speciation events are simultaneous; pseudo- or delayed-cospeciations, where one speciation event happens after the other; host shift, where the parasite acquires a new host (and can subsequently lose the previous host, or undergo speciation); and host-range expansion, in which the parasite acquires a novel host and retains the ability to infect the previous host. When the likelihood of both host and parasite extinction events are added to these models, the variety of outcomes is considerable. De Vienne et al. (2007) shows that host-switch-like events can result in matching trees, and so one could argue that the continental-level signal of tree matching can emerge from numerous host-shift events (although the results of Krasnov et al. 2012a do not support this scenario). Because separating true cospeciation events from all other situations is difficult without time-calibrated trees, de Vienne et al. (2013) suggest that the number of cospeciation events is inflated by cophenetic methods like *PACo* or *ParaFit* (although Lopez-Vaamonde et al. 2001 reported an increased number of cospeciations when accounting for time of speciation events). This reinforces the need for high-quality phylogeny data calibrated for communities in which the interactions are known. Not only will this help us make more robust inference about the evolution of networks (Eklöf & Stouffer 2016), it will also help us leverage information about species interactions embedded within networks to better understand coevolutionary dynamics.

## Acknowledgements

We thank Juan Antonio Balbuena for discussions about the *PACo* method, and members of the Stouffer and Tylianakis groups for comments on an early draft of this manuscript. We thank Scott Nuismer for feedback. We are indebted to Matt Hutchinson and Fernando Cagua for contributions to the code of the paco R package. Funding to TP and DBS was provided by a Marsden Fund Fast-Start grant (UOC-1101) and to DBS by a Rutherford Discovery Fellowship, both administered by the Royal Society of New Zealand.

